# Advances in Thin-Film Graphene Neurotechnology for Chronic Nerve Stimulation and Recording

**DOI:** 10.64898/2026.01.23.701276

**Authors:** Nicola Ria, Bruno Rodriguez-Meana, Eduard Masvidal-Codina, David Andreu, Jose Crugeiras, Lucie William, Anna Graf, Xavier Illa, Georgios A. Katirtsidis, Aina Galceran, David Guiraud, Elena del Corro, Xavier Navarro, Jose A. Garrido

## Abstract

Neurotechnology is being explored for restoring sensorimotor functions after paralysis or amputation, which requires peripheral nerve interfaces that are selective, bidirectional, and chronically stable. Reduced graphene oxide (rGO)-based microelectrodes offer low impedance and a high charge-injection limit; however, long-term in vivo performance has been limited by the durability of encapsulation. Here, we introduce a 10 µm-thick transverse intrafascicular multichannel electrode (TIME) with a hybrid polyimide–Al_2_O_3_ encapsulation engineered to improve fabrication yield, electrode-to-electrode uniformity, and device stability. In vitro, devices maintained near-ideal capacitive behaviour after accelerated ageing (3 months in PBS at 57 °C) and sustained 10^9^ biphasic stimulation pulses without detectable electrochemical degradation. In vivo, the arrays recorded compound nerve action potentials after one month and enabled selective activation of distinct peripheral nerve fibres with comparatively low current thresholds during four months of follow-up, remaining below the device maximum injectable current. Together, these results demonstrate that combining graphene microelectrodes with a thin hybrid encapsulation improves chronic reliability of intraneural thin-film interfaces and helps to close the gap between laboratory prototypes and clinically relevant neuroprosthetic systems.

## Introduction

Interfaces for recording and stimulating peripheral nerves are poised to enable a widening range of therapies and neuroprosthetic systems. Applications span restoration of motor and sensory function in limb loss^1,2^ and neuropathies, neuromodulation of organ function and inflammation via the vagus nerve^3,4^, and treatment of pain^5^ and movement disorders^6^ through targeted peripheral stimulation. Across these indications, translating laboratory success into durable clinical benefits requires neurotechnologies that combine high spatial resolution, selective access to specific fascicles, and stable performance over years. Conventional technologies for nerve interfacing have delivered important advances, but they remain constrained by fundamental trade-offs. Epineural and cuff devices^7,8^ are relatively gentle but often provide limited selectivity. Intraneural longitudinal^9^ (LIFE) or transverse (TIME) devices^10,11^ and Utah-style arrays (USEA) increase selectivity at the cost of more intimate tissue contact and greater mechanical demands. While currently explored clinical technologies have established the feasibility of nerve interfacing, their impact remains limited by invasiveness^12^, mechanical mismatch with tissue^12,13^, and constraints in spatial resolution.

In this context, a new generation of flexible,^14–16^ thin-film technologies promise reduced tissue response^12,17^, a smaller surgical footprint and finer spatial resolution^18^ enabled by miniaturized microelectrodes capable of bidirectional operations^19,20^ (stimulation and recording). However, manufacturing small microelectrodes while maintaining low-noise recordings and deliver sufficient current density to activate fibers reliably remains a major challenge when using clinically approved materials.^21^ To this end, emerging electrode materials and associated fabrication strategies are being explored to raise charge-injection limits and reduce electrode impedance, while preserving the biotic/abiotic stability of the implant. Recent efforts have focused on PEDOT^22^ and related conducting polymers, iridium oxide^23^ (IrOx), and nanocarbon^24^ platforms. Within this landscape, reduced graphene oxide (rGO)-based microelectrodes combine chemical stability, a wide water window and favourable capacitive behaviour^25^ with facile integration on flexible substrates, positioning them as promising candidates for high-density, bidirectional peripheral interfaces. In prior work, we showed in acute settings that rGO microelectrodes exhibit low impedance, a high charge-injection limit (CIL), low stimulation thresholds, and high signal-to-noise ratios (SNR) during neural recording^19,26^.

However, most graphene microelectrode studies have been limited to acute or sub-chronic in vivo use. Translation to chronic use requires advances in thin-film encapsulation to prevent moisture ingress and interlayer delamination while maintaining mechanical compliance. Flexible, biocompatible polymers (PI^18,19^, PDMS^27^, Parylene C^28,29^, SU-8^30^) are widely used, but their relatively high water-vapour transmission rate and their poor interlayer-adhesion can drive degradation under humid conditions^31^ and electromechanical stress^32,33^. To address these limitations, hybrid organic–inorganic stacks have been developed, combining polymer layers with inorganic materials such as HfO_2_^34–36^, SiO_2_^35,37^, SiN_x_^37^, and Al_2_O_3_,^34,38^. These multilayer stacks offer low water-vapour transmission rates (WVTR) and have demonstrated excellent barrier performance in accelerated ageing, although they are often integrated into rigid structures^39^, and their in vivo evaluation remains insufficiently explored^40^.

In this work, we introduce a new generation of transverse intrafascicular multichannel electrode (TIME) technology engineered for chronic peripheral nerve use. We combine (i) a device layout with integrated anchoring features to mitigate displacements, (ii) a hybrid PI–Al_2_O_3_–PI encapsulation that improves moisture-barrier properties and metal–polymer adhesion, and (iii) via-under-electrode routing that isolates rGO residues from conductive tracks. We show that these advances result in high-yield devices with uniform electrochemical performance and robust in vivo function. In bench testing, arrays exhibited excellent electrode-to-electrode uniformity (e.g. CIL and impedance) and no detectable structural change by Raman spectroscopy after accelerated ageing and 10^9^ stimulation pulses. In vivo, rGO microelectrodes achieved selective activation of distinct groups of nerve fibers with thresholds remaining below the maximum injectable current over 120 days, and allowed recording nerve action potentials with robust signal-to-noise ratios. Collectively, our results advance thin-film rGO-based intraneural interfaces towards durable in vivo operation.

## Results

In this study, we designed, fabricated, and tested in vivo a new generation of TIME devices optimized for stable nerve interfacing in chronic applications. The device is a thin-film PI ribbon (10 µm thick, 280 µm wide) comprising 2 arms with eight microelectrodes each (80 µm in diameter, centre-to-centre pitch is 160 µm; Fig. 1a). The structure of each arm was designed with an L-shape layout (Fig. 1a) to prevent longitudinal traction on the implant, which can otherwise displace the electrodes outside the nerve during movement, as previously reported^26^. The layout also incorporates two arrow-like structures (0.55 mm at the base) placed at the centre of the device so that, upon implantation, they rest just outside the nerve to reinforce clamping (Fig. 1b). The edges of the ribbon are also structured to increase friction with surrounding tissue and mitigate device migration.

**Figure 1.**
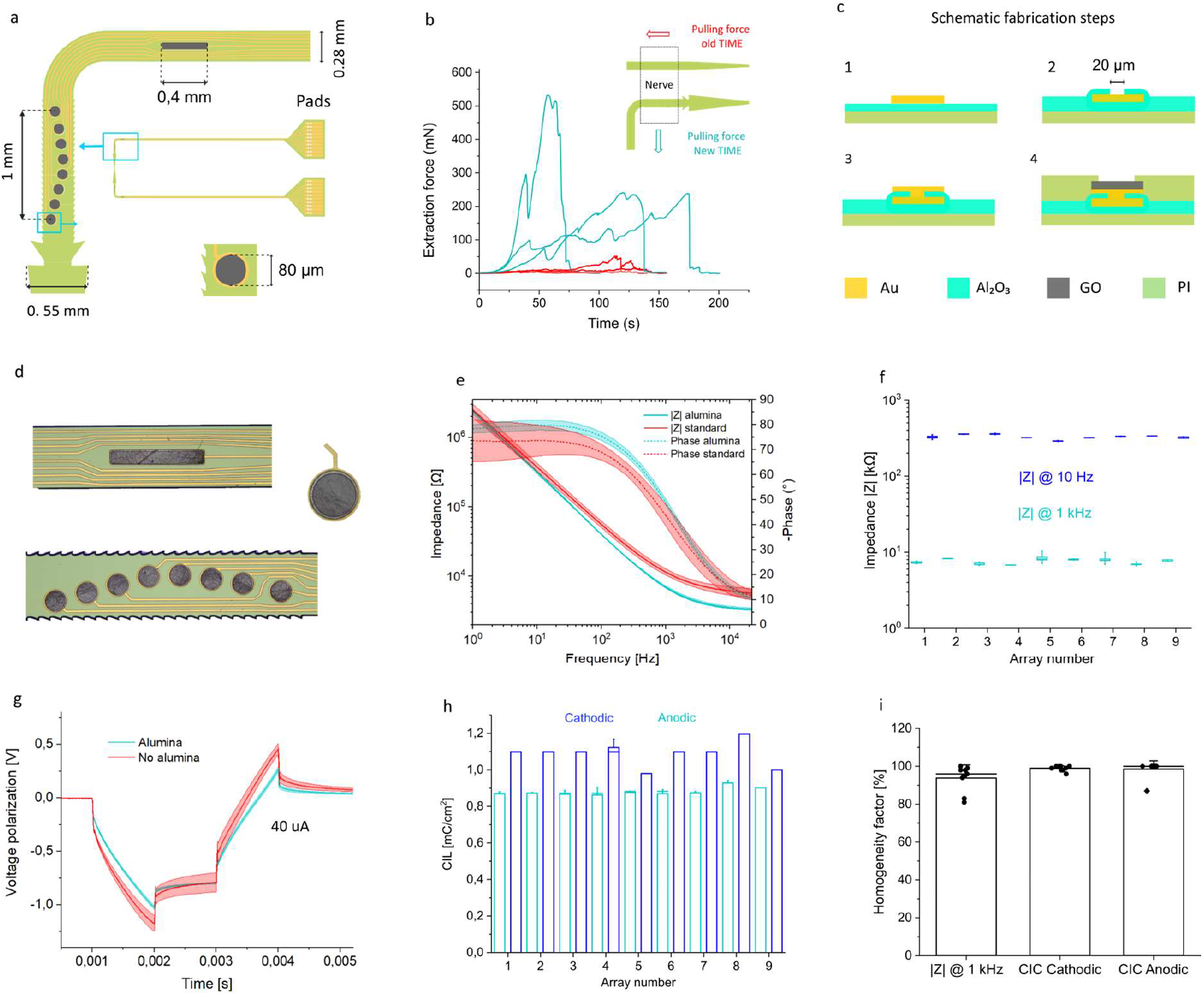
Graphene-based TIME device design, fabrication and electrical performance. **a**, Schematic of the new TIME design: each arm carries an array of eight 80 µm-diameter microelectrodes and a rectangular reference electrode (0.4 mm x 60 µm). The lead has an L-shaped layout and two 0.55 mm anchoring arrow-shaped structures to improve fixation within the nerve. **b**, Extraction force for TIME devices pulled from an explanted sciatic nerve affixed to a petri dish. The graph shows three measurements made with the previous linear TIME design (red) and three with the new design incorporating the anchoring features (light blue). **c**, Cross-sectional fabrication workflow of the device, illustrating the four main process steps. **d**, Optical image of a fabricated device showing the electrode array, with a magnified view of a single microelectrode and the rectangular pad electrode. **e**, Impedance spectra for a PI-PI device (red) and a device with alumina encapsulation (light blue): magnitude (solid line) and phase (dashed line), n=16 electrodes per device. **f**, Impedance magnitude at 10 Hz (blue) and 1 kHz (light blue) for nine arrays (each with eight microelectrodes). Boxplots represent the 25^th^-75^th^ percentiles. **g**, Voltage polarization to biphasic current pulses (1 ms per phase): mean traces ± s.d. (shaded), comparing PI-PI (red) and alumina-encapsulated (light blue) devices (n=16 microelectrodes per type). **h**, Cathodic (blue) and anodic (light blue) charge injection limits for the eight electrodes across nine different arrays. The bar plot represents average values. **i**, Homogeneity factor for impedance magnitude at 1 kHz and for cathodic/anodic charge injection limits across the nine arrays shown in **f** and **h**. The bar plot represents the average value.

To evaluate the clamping capability of this design, three dummy PI devices with the new layout and three with the previous linear layout were inserted^26^ into explanted rat sciatic nerves. Devices were then withdrawn at 100 µm/s with a microcontroller while force was measured with a dynamometer (Fig. S5). The mean extraction force was 337 ± 170 mN for devices with clamping arrows (Fig.1b and S5) versus 26 ± 23 mN for the standard device, more than 10-fold increase.

To improve encapsulation hermeticity and interlayer adhesion, two 50 nm-thick layers of Al_2_O_3_ were incorporated between the metallic tracks and the PI (Fig. 1c). rGO electrodes were contacted to the tracks through vias under the electrodes. Electrode shapes were subsequently defined by photolithography and reactive ion etching (RIE). This patterning step is particularly challenging because rGO residues can remain after RIE; in our design, contacting the metal tracks to the electrodes from below and covering the remaining tracks with a thin, non-conductive layer prevents residual rGO from contacting the gold (Fig. S2a, b). With this change, no inter-electrode short-circuits were observed post-fabrication, resulting in a high fabrication yield (+20%). A photograph of a finalized rGO array, with a magnified view of one electrode, is displayed in Fig. 1d.

We next assessed whether the hybrid PI-Al_2_O_3_ encapsulation affected the performance of the electrodes by conducting electrochemical characterization in phosphate-buffered saline (PBS). Impedance spectra (mean ± standard deviation) from the two arrays of one device, with and without the new encapsulation, showed similar phase and magnitude at low frequency, consistent with near-ideal capacitive behaviour; at higher frequencies, electrodes with the new encapsulation exhibited a lower impedance (Fig. 1e). The mean impedance values calculated for the eight electrodes of nine alumina-encapsulated arrays (n=72) were 329.27 ± 21.64 kΩ at 10 Hz and 7.71 ± 0.98 kΩ at 1 kHz (Fig. 1f), with high uniformity (per-electrode values are shown in Fig. S3). Biphasic current pulses (1 ms) were delivered while recording voltage polarisation at the electrode-electrolyte interface (Fig. 1g). Devices with and without alumina exhibited similar interfacial polarisation, whereas the alumina design showed a smaller ohmic drop, consistent with the reduced high-frequency impedance (Fig. 1e). The charge injection limit (CIL) was determined for both cathodic and anodic pulses across the eight electrodes from nine arrays, yielding average values (n=72) of 1.08 ± 0,06 mC/cm^2^ (cathodic) and 0.88 ± 0.02 (anodic), with high homogeneity in each array (per-electrode values in Fig. S3). Because CIL depends on pulse width, we also evaluated the stimulation protocol used in vivo (biphasic, 50 µs per phase): the maximum safe current averaged ∼600 µA (n=8), corresponding to 0.6 mC/cm^2^ (Fig. S4a,b). Performance homogeneity for impedance and CIL was quantified (refer to Methods for calculation of homogeneity factor, H) across electrodes of the nine arrays (Fig. 1i). The electrodes show a high homogeneity factor for both impedance (H = 93.7 ± 6.9 %, at 1 kHz), and for CIL (cathodic: 98.8 ± 1.3 %; anodic: 98.5 ± 4.3 %).

Overall, these data indicate that this new fabrication method developed to include the alumina encapsulation layer yields high-performance devices with high yield and uniformity.

### In vitro evaluation of microelectrode technology stability

To assess the effect of the hybrid PI-Al_2_O_3_ encapsulation on the devices’ long-term functionality, we subjected them to accelerated ageing followed by prolonged electrical stimulation of the electrodes. To accelerate materials ageing^33,41^, an array was soaked in PBS at 57° C for 3 months (Fig. S6a), corresponding to ∼ 1 year at 37° C by standard acceleration models^34^. To better approximate in vivo use, the same electrodes were then stressed electrically, as current stimulation can accelerate degradation mechanisms^27,42^. We delivered 1 billion biphasic current pulses through 4 electrodes (the maximum simultaneous channels allowed by the software) using the same in vivo protocol (50 µs per phase, 100 µA amplitude, 1 kHz; Fig. 2a), corresponding to a charge density of 0.1 mC/cm^2^, higher than the maximum value approved clinically^33^. While monitoring the electrode polarisation during stimulation (measured for all electrodes using the DC voltage recording capability of the Neuronexus stimulator) revealed that, even with nominally symmetrical pulses, charge accumulated owing to an imbalance in capacitive polarization between anodic and cathodic phases and slow charge balance in between pulses. At high stimulation frequency (1 kHz), the DC offset drifted to −1.6 V within ∼20 s (Fig. 2b), which could initiate faradaic reactions and degradation of the electrode^43^. To mitigate this, we introduced a 1ms passive discharge interval between pulses, which stabilized the baseline near the open circuit potential throughout stimulation^44^ (Fig. 2b). Measurements at neighbouring electrodes at increasing distances from the stimulation site indicated that charge build-up is highly localized around the stimulating site, decaying sharply with distance^45^ (Fig. 2c).

**Figure 2.**
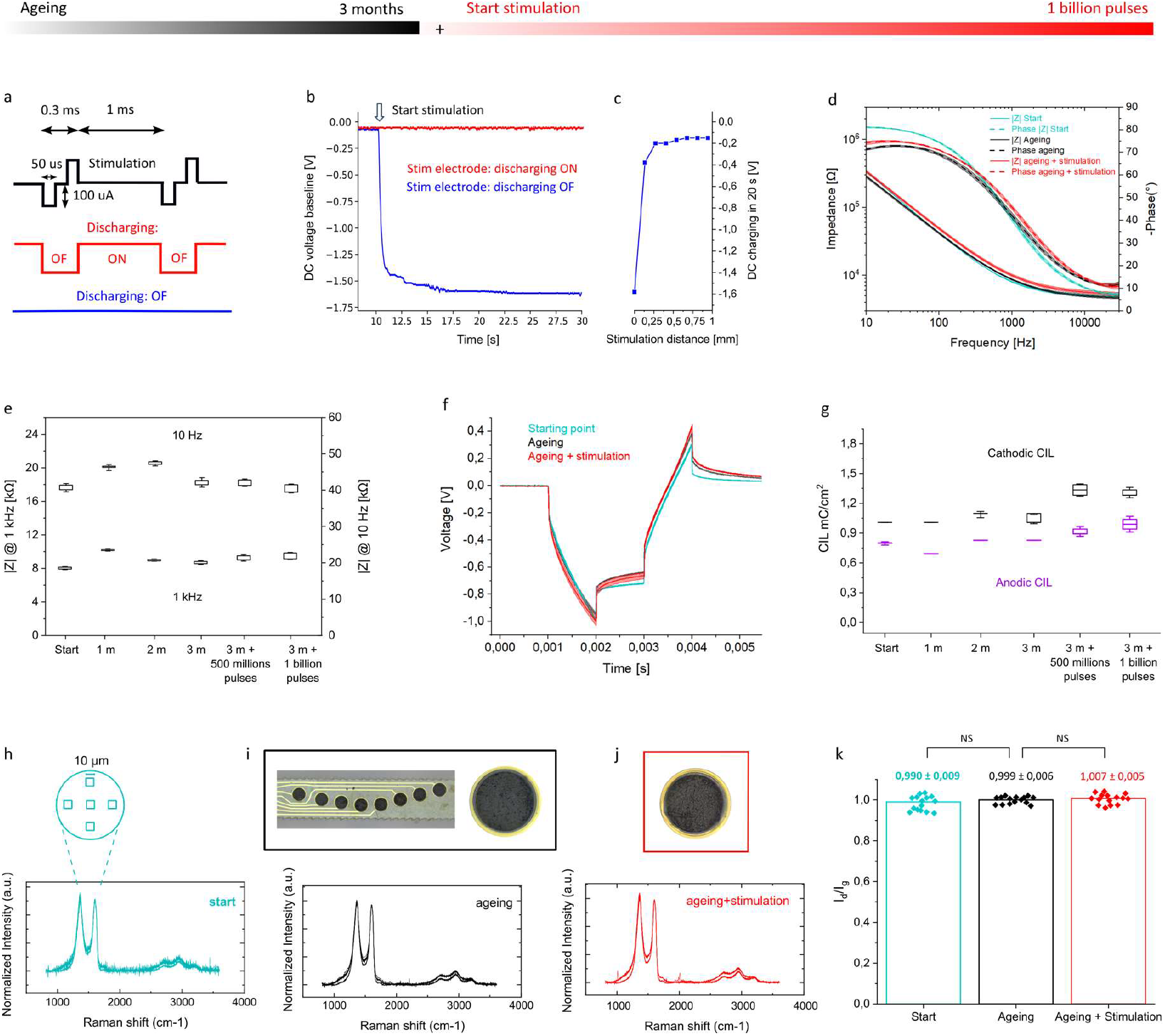
Stability after aging and long-term stimulation. **Top bar**, schematic timeline of the stress test: 3 months of immersion in PBS at 57° C (black), followed by long-term stimulation with 1 billion pulses (red). **a**, Stimulation protocol (black) used for long-term pulsing: biphasic current pulses (50 µs per phase, 100 µA, 1kHz) tested with passive discharge (red) and without discharge (blue). The red bar indicates that, after ageing, the same electrodes undergo up to 10^9^ pulses. **b**, Electrode-electrolyte baseline voltage during stimulation at 1 kHz with one electrode in PBS. Stimulation starts after 10 s and is delivered with (red) and without (blue) inter-pulse discharge. **c**, Baseline voltage reached under the conditions in **b** as a function of the distance to the stimulating electrode (n=8 electrodes). **d**, Impedance spectra (magnitude, solid line; phase, dashed line) for one array at the starting point (light blue, n=8), after 3 months in PBS at 57 °C (black, n=8), and after 10^9^ pulses (red, n=4). **e**, Impedance magnitude at 10 Hz and 1 kHz at the starting point (n=8), after every month of ageing at 57 °C (n=8), and after each 5 x 10^8^ pulses (n=4). Boxplots represent the 25^th^-75^th^ percentiles. **f**, Voltage polarization for biphasic current pulses (1 ms per phase) in the same conditions as **d**; the shadowed area corresponds to standard deviation (s.d.). **g**, Cathodic and anodic charge injection limits under the conditions in **e. h**, (Top) Schematic of the five 10 μm x 10 μm areas sampled for Raman spectroscopy on rGO electrodes. (Bottom) Averaged Raman spectra for 3 different electrodes at the starting point (each averaged over its five areas). **i**, (Top) Optical image of an array (with magnified electrode) after 3 months in PBS at 57 °C. (Bottom) Averaged Raman spectra for the 3 electrodes. **j**, (Top) Optical image of an electrode after 10^9^ pulses. (Bottom) Averaged Raman spectra of the 3 electrodes after long-term stimulation. **k**, Ratio of D-to G-peak intensities from Raman spectra. Points represent the quantification of the ratio across each area of the three electrodes (n=15 per condition). On top, the total weighted mean with standard deviation in the three conditions. The bar plot displays the average value. Statistical tests are described in Methods.

The mean impedance spectra (including standard deviation) for one array at the starting point (n=8), after ageing (n=8), and after 1 billion pluses (n=4) indicated retained electrode functionality (Fig. 2d). Impedance magnitudes at 10 Hz and 1 kHz for different time soaking periods and number of stimulation pulses supported the stability (Fig. 2e) and uniformity of the electrode performance (>94%, S6c). The average voltage polarization (± s.d.) in response to biphasic current pulses of 30 µA amplitude and 1 ms per phase remained consistent in shape and amplitude across all stress conditions (Fig. 2f). The quantification of the anodic and cathodic charge injection limits showed no major change between the starting point (n=8) and monthly ageing time points (n=8), indicating preserved capacitive properties of the electrodes and response uniformity (>95%, Fig. S6b). During extended stimulation (n=4), the CIL increased by ∼30 %, which we attribute to an activation process of the electrodes related to an increase in effective surface area of the porous material, leading to enhanced charge storage and injection limits^25^ (Fig. 2g).

To probe structural changes in rGO electrodes after stress testing, we measured Raman spectra over five 10 μm x 10 μm areas in the electrodes, spanning center and perimeter, to sample a substantial fraction of the electrode surface (Fig. 2h). For three electrodes measured at the starting point, after ageing, and after long-term stimulation, averaged Raman spectra showed no significant changes in peak positions or relative intensities (Fig. 2h-j). The weighted mean (± s.d.) of D/G intensity ratio across the 15 areas in the 3 conditions varied by <1%, suggesting that there is no detectable change in the density or distribution of defects in the rGO electrode material^46^ (Fig. 2k). Furthermore, optical images of the electrodes and the arrays did not evidence water ingress or metal oxidation after the ageing and long-term stimulation treatments (Fig. 2i-j).

Complementing the electrochemical characterization, we examined the encapsulation by SEM on FIB cross-sections (Fig. S7a,b). We compared a PI-Al_2_O_3_-PI stack after 3 months in 57 °C PBS with a standard PI-PI stack soaked for 1 week. In the PI-PI device, a ∼80 nm gap was observed between the first PI layer and the Ti/Au tracks; this gap was not present in the PI-Al_2_O_3_-PI device, indicating improved adhesion at metal-PI interface^47^.

Overall, the electrical and structural analyses indicate that the encapsulation combining PI and Al_2_O_3_ mitigates water ingress and delamination, while the nanoporous rGO electrode material supports stable charge injection limits over prolonged operation and tolerates extended electrical stimulation.

### In vivo chronic functionality for nerve stimulation

The TIME devices were transversely inserted (Fig. S14) into the tibial and peroneal fascicles of the sciatic nerve in rats to evaluate the capability of rGO electrodes to stimulate and record from peripheral nerve fibres over months of implantation. Biphasic current pulses (50 µs per phase) were delivered at increasing amplitude while simultaneously recording compound muscle action potentials (CMAPs) from three target muscles using monopolar needle electrodes. A passive discharge interval was implemented between the pulses to prevent the charge accumulation observed in vitro.

CMAPs recruitment curves were used to quantify neuromuscular activation for a given stimulating electrode. Curve shape at days 30 and 120 remained similar to day 1, with only a small shift towards higher currents (Fig. 3a-c). From the responses produced by the electrodes in the array, we extracted the currents needed to achieve 5%, 30%, and 95% of the maximal CMAP amplitude (Fig. 3d-f). On day 1 (n = 7 rats), the mean current to reach 95% activation was 121 ± 48 µA for gastrocnemius (GM), 182 ± 66 µA for tibialis anterior (TA), and 190 ± 74 µA for plantar (PL) muscles, which is comparable to the values reported for platinum and iridium oxide electrodes of similar size^26,48^. Activation thresholds are expected to drift during chronic implantation owing to the formation of a fibrotic capsule around the device, which lowers tissue conductivity^49^. For 95% activation, we observed mean increases of 21% at day 30 (n=12), and 51% at day 120 (n=12), a modest rise compared to prior reports^26,50–54^. Importantly, current thresholds remained below the maximum injectable current (∼600 µA in PBS; dashed line in Fig. 3d-f) even at day 120, thereby avoiding Faradic reactions that can compromise biocompatibility and long-term functionality^43^.

**Figure 3.**
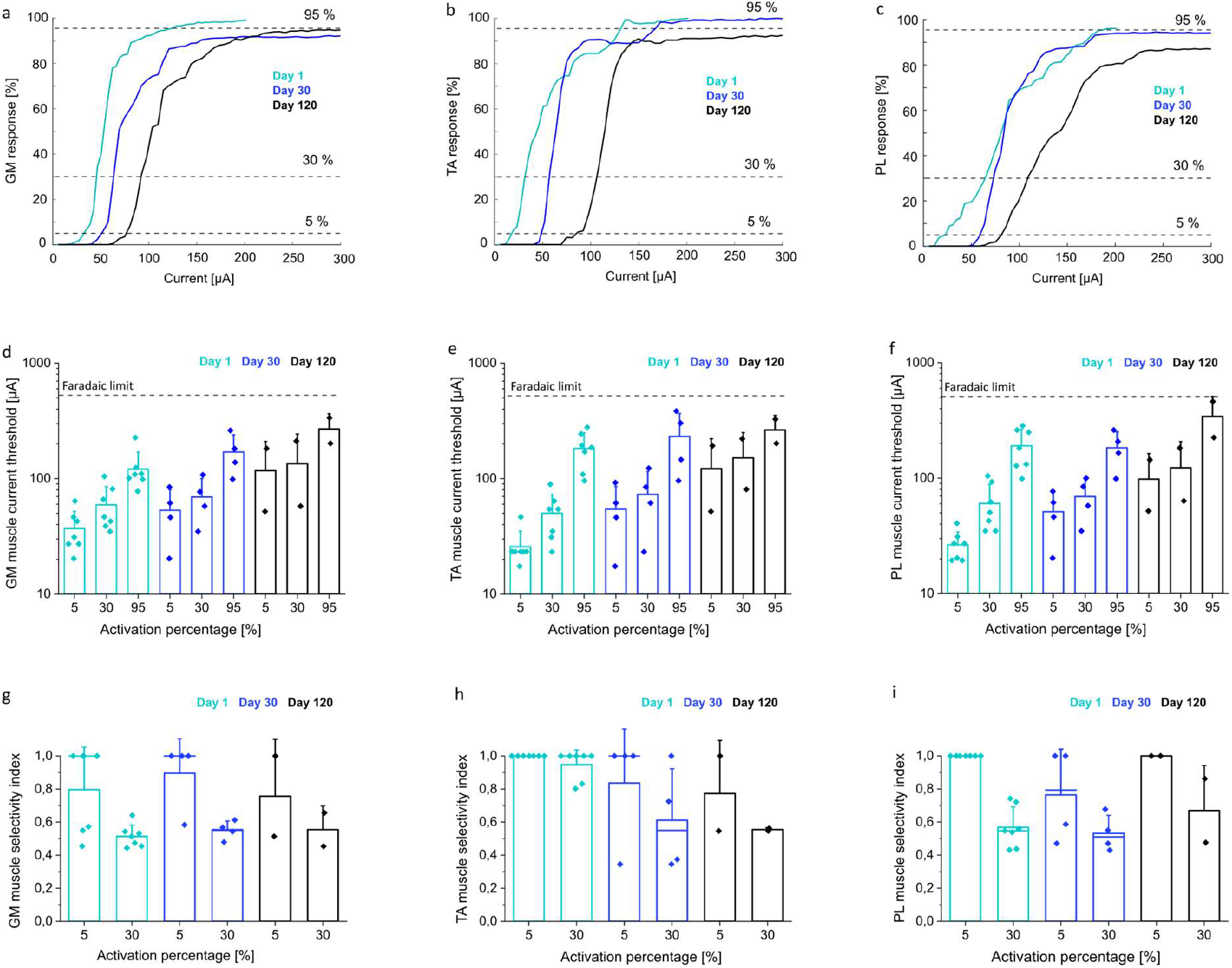
Chronic stimulation of the sciatic nerve. **a-c**, Percentage muscle activation (based on CMAP amplitude) for gastrocnemius (GM; **a**), tibialis anterior (TA; **b**), and plantar (PL, **c**) muscles during stimulation with a single rGO electrode in the sciatic nerve (biphasic current pulses of 50 μs at increasing current intensity). Curves shown for day 1 (light blue), day 30 (blue), and day 120 (black) post-implantation. Dashed horizontal lines mark 5%, 30%, and 95 % of maximal activation. **d-f**, Threshold currents needed to reach 5%, 30%, and 95 % activation for GM (**d**), TA (**e**), and PL (**f**) muscles on day 1, day 30, and day 120. The bar plot displays the average value, and each dot represents the current threshold of one electrode in different implanted animals. The dashed line denotes the maximum injectable current within the water window limits. **g-i**, Selectivity index (SI) at 5% and 30 % activation for GM (g), TA (**h**), and PL (**i**) muscles on day 1, day 30, and day 120. The bar plot displays the average value, and each dot represents the current selectivity of one electrode in different implanted animals.

Previous failures of intraneural stimulation have been attributed to deterioration of the electrodes or the tissue at the interface^53^, displacement of the device outside the nerve^26,54^, or failure of the connection to the stimulator^55^. Maintaining a stable connection with the electronics is very challenging due to the fragility of the contact pads of the device (Fig. 1a), which can cause delamination with time and repeated handling. Here, optical inspection of all explanted devices with alumina (n=7; 30 or 120 days) showed no damage to the gold pads, consistent with the improved adhesion at the metal-PI interface (Fig. S12a). By contrast, devices without alumina (n=9; explanted after 30 days) exhibited partial metal detachment at the pads (Fig. S12b).

Furthermore, no devices (n=7) were found outside the nerve after chronic implantation, which we attribute to the implementation of the new anchoring features. Together, the new mechanical design and alumina encapsulation provided a stable neural interface for at least 4 months. Across the electrodes (n=16) in the two arrays of one representative device, all electrodes could activate at least one muscle up to 95% of maximal CMAP and exceeded 70% in the other two muscles, depending on the relative position of each electrode to the specific nerve fascicle (Fig. S9). Although working electrodes showed high performance, some channels lost functionality to activate the muscles over time (43% non-functional at day 30; 71% at day 120), likely due to limited mechanical stretchability of the device, which is challenged during the leg motion and the repeated access for testing.

Besides the capability to recruit the neuromuscular system, another important functionality is the possibility to activate the nerve fibres with selectivity, particularly in applications aiming to control the coordinated movement of the limbs to restore mobility^56^ or sensation^57^. Dense, small intraneural electrodes delivering focal current can favour selective activation of specific fascicles. A selectivity index (SI, refers to Methods for calculation), which quantifies the activation of one muscle relative to the other two, showed average day 1 values (n=7 for each muscle) of 0.8 and 0.5 for GM at 5% and 30% activation, 1 and 0.95 for TA, and 1 and 0.57 for PL. This index was monitored for each muscle at day 30 (n=4) and 120 (n=2); despite variability, no significant decreases were observed, indicating that the electrodes can stimulate different groups of nerve fibers with good selectivity over time (Fig. 3 g-i).

Overall, despite the limited stretchability of the PI devices, the results indicate that these electrodes can interface stably and effectively with nerves, achieving low activation currents and good selectivity across subfascicles even after long-term implantation.

### In vivo chronic nerve recording

Bidirectional interfacing with the peripheral nervous system, needed to reproduce natural sensations and control complex prosthetic movements, requires both high-resolution stimulation and recording ^58^. Having established chronic stimulation efficacy, we next assessed the recording capability of rGO microelectrodes by measuring compound nerve action potentials (CNAPs) while electrically stimulating the medial plantar (MPN), lateral plantar (LPN), and dorsal peroneal (DPN) nerves of the hind paw with biphasic current pulses (100 μs per phase) delivered via monopolar needles. Representative traces show increasing CNAP amplitude during current ramp for each nerve (Fig. 4a-c).

**Figure 4.**
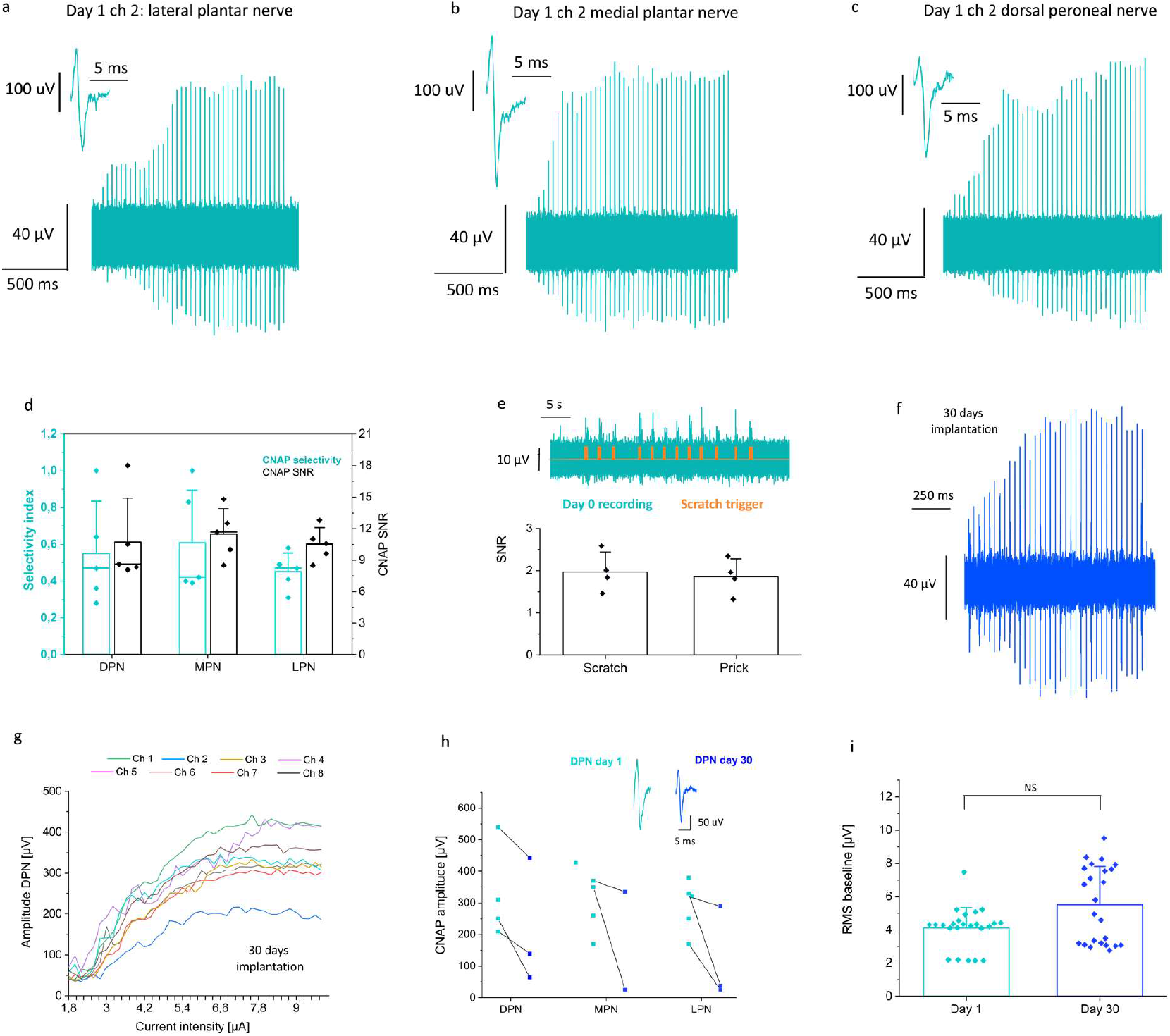
CNAP recordings from the sciatic nerve. **a**, Representative recording, day 1, from a single rGO electrode showing CNAPs elicited by stimulation of the lateral plantar nerve (LPN) with biphasic current pulses (100 μs per phase) at increasing current intensity. Inset, example of CNAP. **b**,**c**, As in **a**, for stimulation of the medial plantar nerve (MPN) and the dorsal peroneal nerve (DPN), respectively. **d**, Selectivity index at 30% activation (light blue) and SNR (black) for MPN, LPN, and DPN across five implants. Bar plots represent the average value. **e**, (Top) Example of ENG recording (band-pass 100-3000 Hz, light blue) recorded from the sciatic nerve during repeated scratch stimuli to the plantar skin (orange ticks mark stimulus times). (Bottom) SNR for scratch and prick stimuli from one electrode in four animals. The bar plot displays the average value. **f**, Representative recording of DPN-evoked CNAPs from one rGO microelectrode at day 30, using the protocol in **a. g**, Recruitment curves of CNAP peak-to-peak amplitude for DPN stimulation recorded with all the electrodes of one array (n=8) at day 30. **h**, (Top) CNAPs recorded at day 1 (light blue) and day 30 (dark blue). (Bottom) Maximum CNAP amplitudes for MPN, LPN, and DPN from one rGO electrode per device at day 1 (light blue, n=4), and day 30 (dark blue, n=3). **i**, RMS baseline noise (1 s window) after stimulation of the three distal nerves at day 1 (light blue, n=24) and day 30 (dark blue, n=24) across eight electrodes per array. Statistical tests are described in Methods.

The peak-to-peak CNAP amplitude exceeded 40 µV for all electrodes in one array (Fig. S8a-c), above the baseline noise (∼30 µV). The average maximum CNAP recorded in 5 different animals were 320 ± 107 µV (MPN), 290 ± 81 µV (LPN), and 287±102 µV (DPN) (Fig. S15). The average signal-to-noise ratio (SNR), calculated as the maximum CNAP amplitude divided by the root-mean-square (RMS) of the full signal, was 11.4 ± 2.4 (MPN), 10.4 ±1.6 (LPN), and 10.7 ± 4.1 (DPN), comparable to high-performance electrode materials^18,19^ (Fig. 4d, black). The dense microelectrode spacing in the array, together with the high SNR achieved with the rGO microelectrodes^18^, enabled discrimination within the sciatic nerve (diameter ≈ 1mm) of responses from a small branch relative the other two, with selectivity indices of 0.6 ± 0.28 (MPN), 0.45 ± 0.1 (LPN), and 0.55 ± 0.28 (DPN) (Fig. 4d, light blue).

To probe the capability to record bursts of single action potentials, we applied brief mechanical stimuli to the plantar surface of the paw. In 4 rats, stimulus-locked bursts of action potentials were detected with mean SNR of 1.97 ± 0.46 (scratch stimuli) and 1.86 ± 0.42 (prick stimuli) (Fig. 4e), in acute as well as in an exemplary recording made at day 30 after implantation (Fig. S10a,b).

Because electrically evoked CNAPs are synchronous, their amplitudes exceeded those from asynchronous mechanical responses by > 10x, allowing reliable chronic detection in more animals (n = 3). Recruitment curves for the eight electrodes of one array showed that all electrodes detected CNAPs > 200 µV after 30 days (Fig. 4g). However, this maximum CNAP amplitude decreased relative to day 1, with reductions ranging from 10% to 93% (Fig. 4h). As with stimulation, recording performance declined over time, in part due to fibrotic encapsulation, which increases resistive tissue around the implant and separates axons from the electrode^59^.

Consistent with this interpretation, impedance spectra from the eight electrodes of one array measured on day 1 and day 30 showed an increase in the magnitude at high frequencies (indicative of higher series/tissue resistance) with similar capacitive behaviour at low frequencies, which is dominated by the electrode properties (Fig. S11). Nevertheless, magnitude remained well below those of a broken channel measured on day 1 (red trace in Fig. S11), indicating that all the electrodes were still functional. This increase in impedance did not generate a significant increase in the recording noise level. The average RMS value (n=24) was 4.1 ± 1.2 at day 1 and 5.5 ± 2.3 at day 30 days (Fig.4i). Despite the noise level was stable, SNR decreased owing to smaller maximal CNAPs (n=3 rats) (Fig. 4h).

Device durability was further evaluated by comparing impedance before implantation and after explantation for two devices used for 30 and 120 days. Few electrodes per device survived the explantation procedure because stretching of the lead during removal broke some metal tracks. In two surviving electrodes, impedance showed no significant change before and after the implantation (Fig. S13).

## Discussion

In this work, we present a new generation of TIME electrode with a design and layer stack engineered for chronic use. The implementation of an L-shaped layout and two arrow-shaped anchoring structures increased the clamping force to the tissue by an order of magnitude. As a result, none of the implanted devices were found outside the nerve after 120 days.

Adding two 50 nm-thick Al_2_O_3_ layers of between the metal tracks and the PI, with a specific stack order, provides a simple route to improve fabrication yield, homogeneity, and robustness in wet environments. Compared with similar multi-layer designs that place tracks and electrodes on different planes separated by μm-thick insulators to compact the layout^60–62^ or to enhance stability,^36,37,39,63^ our approach addresses both aims without significantly increasing the total film thickness (10 μm). Routing the tracks through vias beneath the electrodes also prevents short circuits caused by rGO residues after dry etching, thereby increasing fabrication yield. This strategy is transferable to other applications where sensing sites are defined by dry etching, such as graphene transistors^30,64^ and carbon nanotubes^65^. The absence of fabrication defects and the protection of the tracks during dry etch and following steps yielded highly uniform electrochemical performance (CIL > 98% homogeneity; 1 kHz impedance magnitude > 93% homogeneity).

The hybrid organic/inorganic stack also improves the stability of the lead and the active contacts. Al_2_O_3_ contributes a low WVTR,^66,67^ while PI provides mechanical flexibility^68^. In contrast to stacks that place alumina in the external layers of the film stacking,^34,35^ having this material between PI-PI and PI-metal tracks enhances interfacial adhesion^40^, as suggested by the intact gold pads of explanted devices (Fig. S12); this is a common failure site in neural interfaces^69,70^. After 3 months of accelerated ageing in PBS at 57 °C, cross-sections prepared by FIB and visualized by SEM confirmed good adhesion in the case of alumina-encapsulated devices, in contrast to the delamination observed for devices with no alumina.

The electrochemical performance of electrodes tested in vitro showed no significant change after combined accelerated ageing and long-term stimulation (10^9^ pulses). Incorporating a passive discharge interval in the stimulating protocol, in vitro and in vivo, prevented charge build-up and drift of the potential beyond rGO’s safe water window. The combination of balanced-charge injection protocols and the intrinsic electrical properties of rGO effectively mitigated structural damage in the microelectrodes; Raman spectroscopy showed <1% change in the D/G intensity ratio. Optical inspection of the leads revealed no water ingress between the PI layers and no metal corrosion, consistent with the low moisture permeability^40^ of Al_2_O_3_ and the chemical resistance of PI^68,71^.

In vivo, TIME devices overcame previous limitations in recording sensory-evoked nerve activity and CNAPs after 30 days of implantation. Signal amplitude decreased over time, which we partially attribute to fibrotic capsule formation^26^; degradation of the electrodes would be expected to increase the noise levels of the recording, which was not observed. The increase in impedance at high frequencies after 30 days is consistent with a higher resistance generated by the fibrotic tissue. Notably, the explanted working electrodes exhibited an impedance comparable to values measured before implantation. Anti-inflammatory surface chemistries (e.g. dexamethasone-functionalized PI) have reduced fibrotic encapsulation around implanted devices^72^, which may further improve long-term stability when combined with our approach.

For chronic stimulation, nerve fibers were activated with high selectivity (SI > 0.5) and with current thresholds slightly lower than those reported for IrOx electrodes of similar size.^48,54^ Electrodes were able to activate all three muscles even after 120 days of implantation, with current thresholds slightly higher due to the fibrotic encapsulation (e.g. +51% at 95% muscle activation after 120 days) but below the safety current limit of the electrodes (∼600 µA). The capability to chronically activate muscles while maintaining the current injection below the safe charge-injection limits of the electrode material is essential to avoid Faradaic reactions at the electrode–tissue interface that compromise long-term functionality. Here, this was enabled by the high CIL of rGO microelectrodes together with improved device alumina-based encapsulation, which reduces water ingress and prevents inter-track short circuits.

The principal limitation of this study is the low mechanical stretchability of the lead. Leg motion and repeated access can induce mechanical loads that fracture metal tracks, leading to the loss of some electrodes over time; assessing alumina integrity under repeated bending also remains important. Future work should explore stretchable yet hermetic neuroelectronic architectures for nerve applications^4,73,74^, for example serpentine/mesh conductors or elastomeric encapsulation with embedded inorganic barriers.

Overall, these advances increase the maturity of thin-film intraneural interfaces by improving homogeneity, stability, and fabrication yield. In vitro and in vivo results demonstrate substantial gains over previous generations, narrowing the gap between laboratory prototypes and clinical translation.

## Methods

### Fabrication of TIME devices

The fabrication started by spin coating a 7.5 µm of PI (PI-2611, HD MicroSystems) onto a Si/SiO_2_ (5000 µm, 285 nm) wafer. The PI was then baked at 350°C in a nitrogen-rich atmosphere, and a 50 nm layer of Al_2_O_3_ was deposited at 200°C using plasma-assisted atomic layer deposition (ALD) (Picosun, R200 Advanced). Then, 20/200 nm thick Ti/Au metal tracks were defined with photolithography using an image reversal photoresist (AZ5214E, Merck) and e-beam evaporation (Leybold Univex 400) in a lift-off process (Fig. S1). A second 50 nm layer of Al_2_O_3_ was deposited on top, and with photolithography using a positive photoresist (HiPR 6512, FujiFilm) and reactive ion etching (RIE), vias were opened at the electrode and pad positions. RIE was performed with a HF power of 50 W and ICP of 800 W for 9 minutes with 20 sccm of CHF_3_ and 30 sccm of CF_4_. A second 200 nm layer of gold was evaporated on top to contact the metals below the vias. A GO membrane was obtained by filtration of an aqueous solution of dispersed graphene oxide flakes (Global Graphene Group) and then transferred onto the previously prepared wafer. The membrane was partially reduced hydrothermally in an autoclave at 134° C for three hours. The electrode areas were defined by using an aluminium hard mask patterned using optical lithography (nLOF 2070, Microchemicals) and lift-off. Then, the GO membrane was etched using RIE with 2000 W ICP and 40 sccm of O2 and 20 sccm of Ar. Both the aluminium on top of the electrode and the electrodes themselves were used as a mask to wet etch the gold from all the wafer except below the electrodes. Afterwards, a second layer (2.5 µm thick) of PI was spin-coated on top. A series of photolithography (AZ 10XT, Microchemicals) and RIE processes were used to open the pads, electrodes, and structure sides. RIE was performed using a HF power of 40 W, ICP power of 1200 W, with 60 sccm of O_2_ and 50 sccm of Ar flow for 4.5 min to etch the top PI layers that opened the electrodes. For the pads and the structure, a second RIE process was applied to open the intermediate alumina using the conditions mentioned before. To open the device structure, an extra RIE process was performed to etch the bottom PI layer. Finally, the aluminium on the electrodes was chemically etched using an aqueous etching solution of orthophosphoric acid, nitric acid, and deionized water (74% v/v H_3_PO_4_, 2.5% v/v HNO_3_, 2.5% H_2_O).

### Electrochemical characterization

#### Set-up

The electrochemical characterization of the electrode was performed as previously reported^18^; here below, we provide a brief summary. A potentiostat (Metrohm Autolab PGSTAT128N) was used in a three-electrode configuration. A platinum wire (Alfa Aesar, 45093), providing sufficient active area, was used as the counter-electrode, and an Ag/AgCl electrode (FlexRef, WPI) as the reference electrode. The TIME device was connected to a 10-contact Zero Insertion Force (ZIF) Molex connector (Mouser Electronics, 0,5 mm pitch), which was attached to a custom small printed circuit board (PCB) (WURTH ELEKTRONIK). A 20-channel header connector (DigiKey, 2.54 mm pitch) connected the PCB to the potentiostat. The device was immersed in a solution prepared by dissolving one PBS tablet (Sigma-Aldrich, P4417) in 200 ml of distilled water to reach the final concentration of 10 mM phosphate buffer, 137 mM NaCl, and 2.7 mM KCl at pH 7.4.

#### Electrochemical activation

Before in vitro electrochemical evaluation, the electrodes underwent an electrochemical activation process consisting of 70,000 biphasic rectangular current pulses (cathodic first with delay) of 1 ms pulse width, ramping from 5 to 40 µA at 100 Hz. Electrochemical characterization in PBS involved measuring impedance, cyclic voltammetry, and current pulses to evaluate functionality, ensuring no broken or electrically shorted electrodes.

#### Impedance spectroscopy

was measured using the above described three-electrode setup by applying a 10 mV sinusoidal voltage excitation over a frequency range from 1 Hz to 100 kHz. Electrodes with impedance between 2 and 20 kΩ at 1 kHz were considered functional.

#### Chronopotentiometry

Biphasic current pulses were injected to individual electrodes while the interfacial potential was measured against the reference. For current pulses of 1 ms, the CIL is determined by looking at the cathodic and anodic voltage shifts. These values were determined for each current level by visually distinguishing the voltage shift across the electrode-PBS interface from the linear voltage produced by the resistive ohmic drop.^43^ To determine the charge injection limit, voltage shift values were plotted against the current for both the cathodic and anodic parts of the pulse at increasing current levels. From the linear fit of these points, we extrapolated the current values at 0.8 V and −0.9 V, representing the maximum currents safely injected within the rGO water window.

#### Homogeneity factor (H)

This parameter was calculated from the mean of the impedance magnitude at 1 kHz and the cathodic and anodic charge injection limits, for the electrodes of each array, and their standard deviation, by using the equation H = 100 (1−STD/Mean)

### Complementary characterization

#### Raman spectroscopy

Raman scattering spectroscopy was performed at five evenly spaced regions (10 × 10 µm^2^, ∼1 point/µm^2^) of three electrodes for each condition (‘Initial’, ‘After ageing’, ‘After ageing + 1B pulses’). A WITec alpha300 R spectrometer was used for all the measurements, with a 600 gr/nm, and a laser excitation wavelength of 488 nm with a 50x objective over 10 s exposures and three accumulations per point. Each Raman map was acquired with ControlSIX software, analysed by fitting the bands of interest (D band ∼1350 cm^-1^ and G band ∼1590 cm^-1^) to 2 Lorentzian peaks, after performing a polynomial background subtraction, and averaged them for each condition. The amplitude intensity ratio between D and G bands was calculated to monitor changes in the defect density throughout the ageing process. In Fig. 2K, the weighted mean is calculated as 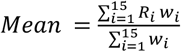with 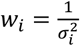 and σ the STD.

### In vitro test

#### Charge building up

Biphasic current pulses (50 µs per phase, 100 µA, and 1 kHz) were applied through the first electrode of the array of one device in 10 mM PBS, using the Neuronexus stimulator (X-Series XDAQ Core Family neurostimulator and a XDAQ Smartbox Pro X3SR head stage which carries an Intan RHS 2000 chip). The stimulation was performed with and without 1 ms discharging in between the pulses. From the DC recording of all the electrodes, one point was extracted between the pulses every 12 ms to plot the voltage baseline versus time. This baseline was smoothened from noise fluctuation with the Savitzky-Golay filter with a second-order polynomial fit in segments of 15 points.

#### Long-term stimulation

To evaluate the long-term stability of the electrodes, biphasic current pulses (50 µs per phase, 100 µA, 1 kHz) were delivered using the StimRG stimulator (Neurinnov, France). Stimulation was applied simultaneously through four rGO electrodes, which is the maximum number supported by the software. The setup used a multipolar push–pull configuration: the four rGO electrodes injected pulses in phase, while a platinum wire delivered pulses of the opposite phase. A second platinum wire was used as the counter electrode for the current return path. Charge-balancing, integrated in the stimulator, was applied between consecutive pulses to prevent charge accumulation at the electrode–tissue interface. Electrochemical stability was monitored by measuring impedance and chronopotentiometry with a potentiostat after every 500 million pulses.

#### Ageing test

Accelerated ageing test were performed to evaluate the lifetime of the hybrid PI-Al_2_O_3_ encapsulation. The devices were immersed in a beaker with 10 mM PBS, hermetically sealed with parafilm, and placed in a heating station at 57°C. By increasing the temperature, it is expected to accelerate the degradation of the polymers used in biomedical applications^75^, with an expected accelerated aging factor of 4, calculated using Arrhenius′ law as reported in previous studies^41^. Devices were removed from the beaker and characterized each month, with the standard setup.

#### Measurement of insertion and buckling forces

Dummy devices of only PI, of both standard (n=3) and improved (n=3) designs, were inserted in the sciatic nerve of rats, with the same procedure as the regular implantation in vivo (refer to in vivo surgery for more details). The nerve was explanted by cutting proximally and distally to the dummy devices. The nerve stumps were fixed with tape on a petri dish, and each device was clamped to the arm controlled by the micro driver (Zwick Roell Z 0.5). The extraction force was measured using a load force cell (Zwick Roell, nominal force 50 N) by pulling out the devices from the nerve at 100 µm/s until complete extraction.

### In vivo procedures

#### Ethics

Animal experiments were conducted in accordance with protocols approved by the Ethical Committee of the Universitat Autònoma de Barcelona and in compliance with the European Communities Council Directive 2010/63/EU. Animals were housed at 22 ± 2 °C under a 12 h light/12 h dark cycle with food and water available *ad libitum*. Appropriate measures were implemented to minimize pain and discomfort during surgical procedures and throughout the postoperative period.

#### Implantation procedures

Surgical procedures were carried out under anaesthesia using ketamine/xylazine (90/10 mg/kg, i.p.) in female Sprague–Dawley rats (250–300 g, ∼ 18 weeks old). Prior to implantation, the TIME device was sterilized with ethanol 70% and carefully folded at the central point so that the two arms were perfectly overlapping. Implantation was performed using a straight needle attached to a 10–0 loop thread^26,48^ (STC-6, Ethicon), which guided the device transversally through the insertion across the tibial and peroneal branches of the sciatic nerve (Fig. S14a-c). The tibial fascicle innervates both the gastrocnemius (GM) and plantar interosseous (PL) muscles, whereas the peroneal fascicle innervates the tibialis anterior (TA) muscle. All manipulations of the nerve and devices were done using fine forceps under a dissection microscope to ensure precise positioning of the microelectrodes within the nerve fascicles; the ending clamp arrow was left just outside of the nerve to avoid possible pulling of the device. After completing the nerve stimulation and recording protocols, as detailed below, the ribbon of the device was routed through the muscle incision and the pad section positioned subcutaneously to allow easy access, and enclosed in a plastic envelope sealed with Kwik-Cast to protect it from damage and fibrotic encapsulation (Fig. S14d). The muscle layer was closed with silk sutures, and the skin incision with small surgical staples. In chronic experiments, the skin and the envelop were reopened, and the pad area was gently cleaned and reconnected to a ZIF connector to perform the required experiments. Throughout the procedure, the animal body temperature was maintained using a heating pad.

After completing the experiments, the devices were explanted. The arms of the device were carefully followed toward the nerve, gently removing any surrounding tissue. The nerve segment containing the implanted device was harvested and immersed in a collagenase solution to dissolve the tissue and release the electrode without applying additional mechanical force to the device.

#### Impedance measurement in vivo

The device contact pad was connected (with the same connector and PCB used for the electrochemical characterization in vitro) to a PalmSens 4 potentiostat. The impedance was measured at each of the microelectrodes, using a stainless-steel needle electrode placed in the adjacent muscle as the counter electrode, and a reference electrode positioned at the base of the tail. The impedance was recorded across a frequency range from 50 kHz to 1 Hz in 47 steps.

#### StimRG stimulator

The stimulator was specifically developed to drive the new TIME electrode. The core technology is based on previous developments.^76,77^ The stimulator has 16 current-controlled outputs, among which at most 4 anodes and 4 cathodes are driven simultaneously to provide multipolar stimulation. The stimulation main features include: i) 4 balanced biphasic with a 100 µs interphase waveforms (rectangular /ramp up, (a)symmetric), ii) 10-500 µs pulse width with 2 µs step, iii) 1-6000 µA (sum on up to 4 simultaneous outputs), with a minimal step of 1 µA. To ensure safe and reproducible stimulation, the stimulator operates at 20 V, capable of driving high-impedance loads with very precise current control and passive charge balancing, thanks to a high-precision ASIC design.

#### Stimulation procedure

First, the maximal muscle response was determined by delivering rectangular pulses (100 μs duration, 1–10 mA) using a Grass S44 stimulator with a PSIU6 isolation unit and two small needles positioned near the sciatic nerve. The evoked CMAPs were recorded from PL, GM, and TA muscles using small stainless steel needle electrodes (13 mm length, 0.4 mm diameter; A-03-14BEP, Bionic) (Navarro, 2016). The active electrode was inserted in the muscle belly, while the reference electrode was placed near the distal tendon. Electromyographic signals were amplified (P511AC amplifiers, Grass), band-pass filtered (3 Hz–3 kHz), and digitized at 20 kHz using a PowerLab16SP system (ADInstruments).

Subsequently, nerve stimulation through each of the microelectrodes in the implanted TIME device was performed by applying two series of 52 biphasic current pulses (50 μs per phase) at 3 Hz using the StimRG stimulator (Neurinnov, France). The current intensity was progressively increased from 0 µA up to the threshold that evoked the maximal muscle response or to a maximum of 600 µA, using each microelectrode as cathode, and the rectangular rGO electrode (Fig. 1a) or a small needle in contact with the sciatic nerve as anode. A counter electrode was positioned at the base of the tail. The CMAP recordings were made as above.

#### Recording procedure

The CNAPs were recorded from each microelectrode, following electrical stimulation of three distal nerves in the hind paw: the medial plantar nerve (MPN), the lateral plantar nerve (LPN), and the dorsal peroneal nerve (DPN). In acute experiments, the reference electrode was the rectangular rGO electrode (Fig. 1a), whereas in chronic experiments, it was a small needle placed near the sciatic nerve. In both cases, the ground electrode was a small needle inserted into the skin at the mid-thigh. Stimulation was delivered using 50 biphasic rectangular pulses (100 μs duration, 0 to 10 mA; DS4 Stimulator, Digitimer) in bipolar configuration via two small needle electrodes inserted near each nerve, into the medial, lateral, or dorso-medial regions of the paw.

In a second protocol, evoked electroneurography (ENG) activity was recorded from the rGO microelectrodes implanted in the sciatic nerve in response to mechanical stimulation of the paw. The configuration of recording electrodes was the same as in the CNAPs recording. Brief scratches were applied with a thin probe, and pricks were delivered using a blunt 27-G needle. These stimuli were synchronized using a push-button trigger (ADInstruments) to facilitate analysis. The evoked CNAPs and ENG signals were amplified (P511AC, Grass), band-pass filtered (300 Hz–10 kHz), processed through a noise eliminator (HumBug, Quest Scientific), digitized at 20 kHz, and recorded using LabChart software (PowerLab System, ADInstruments).

### In vivo data analysis

#### Analysis for stimulation experiments

The CMAP recordings were exported and analysed using custom MATLAB scripts (MathWorks, R2023b). The CMAP waves were identified within predefined 10,3 ms time windows, corresponding to each stimulation event. Since stimulation was delivered twice for each microelectrode, the peak-to-peak CMAP amplitude of the two repeated trials was measured and averaged to enhance reliability. These values were normalized as a percentage of the maximal CMAP of each muscle and animal, and used to generate recruitment curves, illustrating the relationship between stimulation intensity and neuromuscular activation. To assess the selectivity of stimulation, the selectivity index was calculated for each electrode configuration^78^. This index is computed as the ratio of the normalized response of an individual muscle to the sum of responses across all monitored muscles. Recruitment threshold and selectivity were evaluated at three relative activation levels: 5%, 30%, and 95% of the maximum observed CMAP amplitude. These levels were chosen to represent distinct functional activation scenarios. A 5% threshold represents the minimal activation of the muscle, considered to avoid potential crosstalk effect; the 30% threshold corresponds to typical muscle activity needed for everyday tasks (Paternostro-Sluga 2008), and the 95% threshold reflects near-maximal muscle activation.

#### Analysis for recording experiments

Electrophysiological signals recorded in vivo were processed using Python packages (Numpy 1.21.5, Neo 0.11.1, pandas, seaborn 0.11.2, Matplotlib 3.5.1, Quantities 0.13.0, Elephant 0.10.0), and the custom library PhyREC 0.6.5. Origin 2018 was used for scatter plots, box and bar charts and statistical analysis.

The recorded signal was first cleaned from voltage artefacts produced by the stimulation pulses. A voltage threshold of 0.4 V was implemented in the trigger channel to detect the time point of the artefact events. Then the signal in the real recording channel was set to 0 in a time window (from 1.1 to 2.1 ms) around the detection, depending on the specific experiment and current level. A refractory period of 15 ms was employed to ensure that the same event was not detected twice. A second threshold was employed to detect the time point of the actual CNAP, and the peak-to-peak amplitude was measured. In the ENG recording, the Root Mean Square (RMS) was calculated in 1 second recording pre-filtered for noise-cleaning (50 Hz and harmonics), in a frequency range of 200-2000 Hz, and then computing the RMS of the squared signal values. The SNR was calculated as the ratio between the maximum RMS detected by each microelectrode during each mechanical stimulus, and the respective RMS of 1 1-second recording baseline before the stimulation.

#### Statistics

For each data set, a normality test was first performed to evaluate if the data followed a normal distribution (P > 0.05) or not (P ≤ 0.05). In the first case, we quantified the statistical significance with the paired t-test for groups of the same size, and Welch’s t-test for groups of different size. For data not following a normal distribution, we used the Mann-Whitney U Test. Significance levels were indicated as follows: NS (not significant, P > 0,05), * (P ≤ 0,05), ** (P ≤ 0,01), *** (P ≤ 0,001), **** (P ≤ 0,0001). All the statistical tests were two-sided. The bar and box plots are expressed with the median line and STD whiskers. All the tests were performed with Origin 2018 software.

Figure 2k, NS p = 0,085 start-aging; NS p = 0,34 ageing–ageing and stimulation, in the paired T-test, Figure 4i, NS p = 0,081 in the Mann-Whitney U-Test. i, NS: p = 0.14 in the Mann-Whitney U-Test.

## Data availability

All the data in the main text has been deposited at Supplementary experimental data that support the figures and other findings of this study can be obtained by contacting the corresponding authors. Authors can make data available on request, agreeing on the data formats needed.

## Code availability

The custom code developed for neurophysiological analysis has been deposited on GitHub and CORA.

## Acknowledgments

This research was funded by FLAG-ERA JTC 2021 project RESCUEGRAPH, by the Agencia Estatal de Investigación of Spain projects PCI2021-122075-2A, PCI2021-122095-2A and CNS2023-144492; and by CIBERNED (grant CB06/05/1105), CIBER-BBN (CB06/01/0049) and TERAV (RD21/0017/0008) funds from the Instituto de Salud Carlos III of Spain, co-funded by European Union (NextGenerationEU, Recovery, Transformation and Resilience Plan). ICN2 is supported by the Severo Ochoa Centers of Excellence programme (Grant CEX2021-001214-S), funded by MCIN/AEI/10.13039.501100011033, and by the CERCA Programme of Generalitat de Catalunya; and by the European Union NextGenerationEU/PRTR. IMB-CNM is supported by the María de Maeztu Units of Excellence programme (Grant CEX2023-001397-M) funded by MCIN/AEI/10.13039.501100011033. N.R. acknowledges fellowship PRE2020-093708 funded by MCIU/AEI /10.13039/501100011033 and by “ESF Investing in your future”. This publication has received funding from the European Union’s Horizon Europe research and innovation programme under grant agreement No 101158723-GAIA and No 101070865 (MINIGRAPH). A.G. thanks La Caixa Foundation Inphinit fellowship LCF/BQ/DI22/11940031. This work has made use of the Spanish ICTS Network MICRONANOFABS, partially supported by MICINN and the ICTS NANBIOSIS, specifically by the Micro-NanoTechnology Unit U8 of the CIBER-BBN The authors acknowledge the assistance of Isidoro Soto from Zwick Roell for performing the force measurements. The authors acknowledge the assistance of Milan Demarcq for designing the StimRG.

## Author contributions

N.R. worked on design and testing of the fabrication protocol with the hybrid PI-Al_2_O_3_ encapsulation; design, fabrication, and characterization of the devices, in vivo recording analysis, and preparation of the manuscript. B.R-M, J.C. and L.W performed the electrophysiological in vivo studies and data analysis. E.M.C. contributed to electrode design, device characterization, and data analysis. A.G. performed the Raman study. X.I. contributed to the fabrication of the devices. G.A.K. contributed to the hybrid PI-Al_2_O_3_ encapsulation technology. A.G. contributed to device characterization. D.G. and D.A. supervised the stimulator (hardware and software) specifications and development. D.G. contributed to the electrical stimulation protocol design. X.N. supervised the electrophysiological studies. J.A.G. supervised the development of the technology and the arrays characterization. X.N. and J.A.G. coordinated the funding projects. All authors contributed to the text of the manuscript.

