## Supplementary information for "Advances in Thin-Film Graphene Neurotechnology for Chronic Nerve Stimulation and Recording"

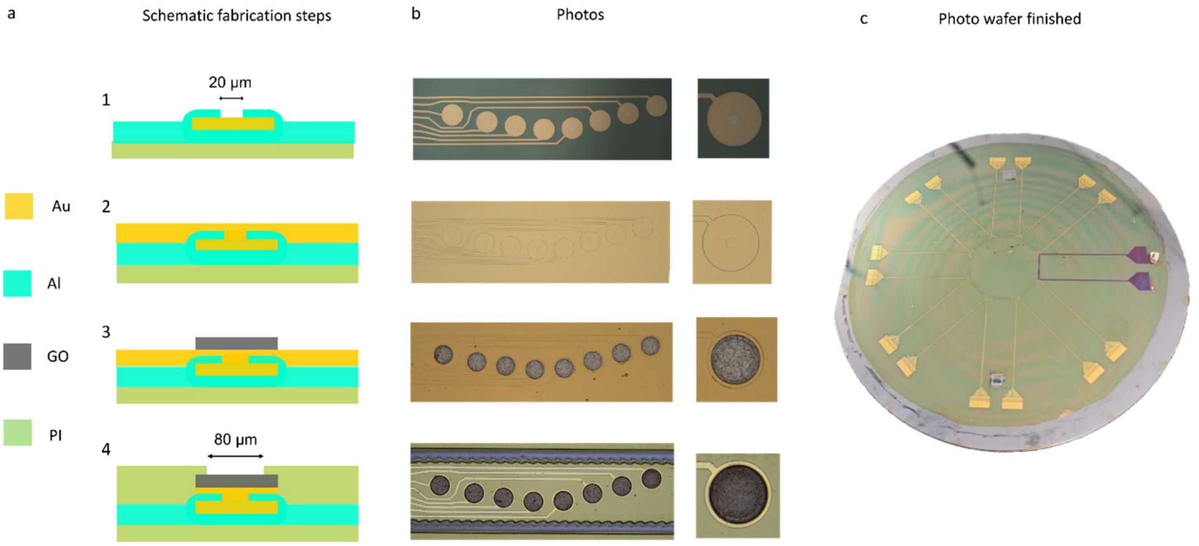


**Figure S1 | Fabrication process. a,** Schematic of the four main steps of the fabrication protocol in sequence: 1. opening of vias in the second alumina layer, 2. gold evaporation, 3. GO transfer, and electrode area definition, 4. deposition of the second PI layer with an opening at the electrode site. **b,** Optical images of the fabrication steps shown in **a**. **c,** Photograph of a completed wafer with one device delaminated.

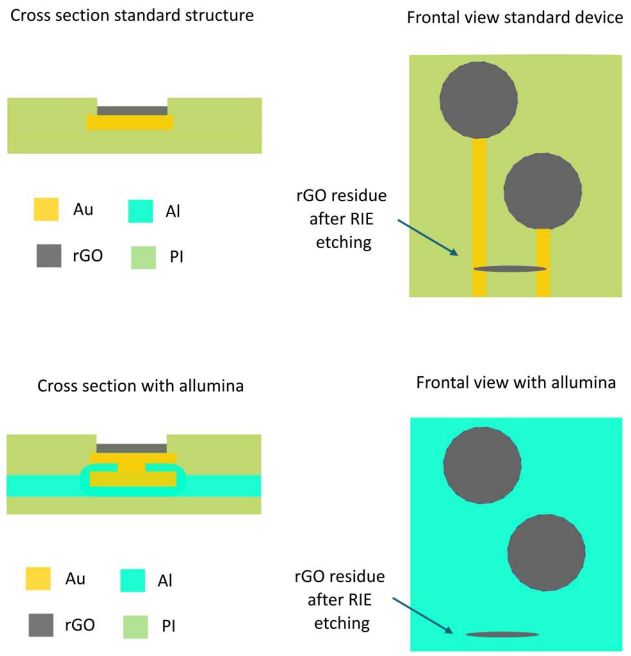


**Figure S2 | Impact of rGO residues in the standard and the new encapsulation procedures. a**, Schematic of the cross-section and frontal view of the standard fabrication. The rGO residues contact the gold, creating electrical shorts. **b**, Schematic of the cross-section and frontal view of the new structure. The rGO residues don't contact the gold tracks, since they are packed between two alumina layers.

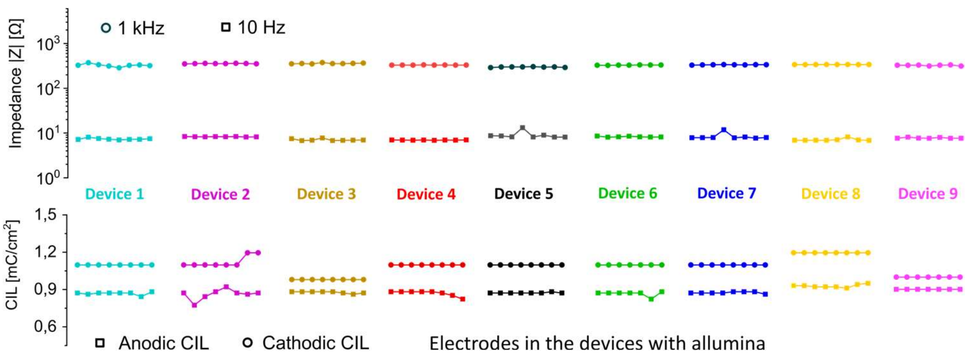


**Figure S3 | Performance and homogeneity.** **a**, Impedance magnitude values at 1 kHz and 10 Hz for all electrodes of 9 different arrays. **b**, Cathodic and anodic charge injection limits for the same electrodes shown in **a**.

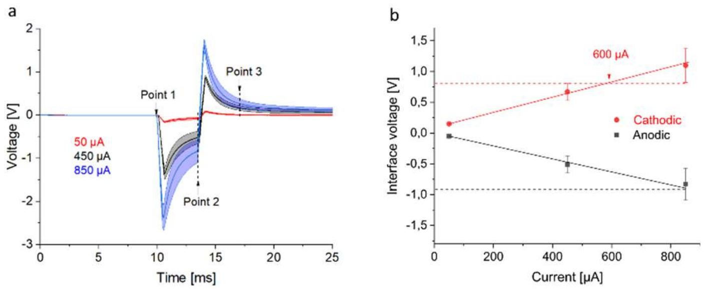


**Figure S4 | Calculation of charge injection limit for short pulses. a,** Average voltage polarization (solid lines), with shadowed area for standard deviation (STD) (n=8 electrodes), of biphasic current pulses of 50 µs in pulse width and 3 ms in inter pulse interval, for 50 (red), 450 (black), and 850 (blue) µA current amplitude. **b**, average cathodic (point 2 – point 1) and anodic (point 3 – point 2) at the electrode-PBS interface with STD (n=8) for the pulses of subpanel a. The dashed line represents the safe voltage limits for the anodic (black) and cathodic (blue) polarizations, from which the maximum current injectable (600 µA) is extracted.

In the case of small pulses of 50 µs, since it was not possible to distinguish the voltage shift from the ohmic drop, a long interpulse interval of 3 ms was applied, to extract the maximum charge injection by considering the difference between the voltage baseline at the beginning and end (3 ms after) of the pulse (Figure S4 a, point 1 and 3) and the voltage recovered at the end the interpulse interval (Figure S4a, point 2). In particular, the cathodic voltage was calculated as $V_{point2} + V_{point1}$, while the anodic $V_{point3} + V_{point2}$. In Figure S5b, the error bars are calculated from the standard deviation (σ) of the three voltage points $V_{point1} \pm \sigma_1$, $V_{point2} \pm \sigma_2$, $V_{point3} \pm \sigma_3$, as $E_{cathodic}=\sqrt{\sigma_1^2 + \sigma_2^2}$ and $E_{anodic}=\sqrt{\sigma_2^2 + \sigma_3^2}$.

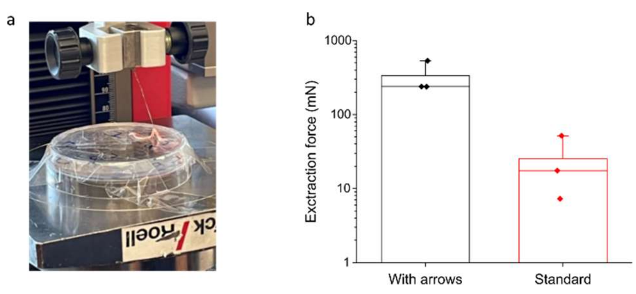


**Figure S5 | Clamping force**. **a**, Set-up for measuring the force needed to extract a device from an explanted nerve from a beaker. The nerve is attached to the beaker, while the device is pulled outside the nerve with a micro driver. **b**, The extraction of the maximum force needed to detach the PI device, for 3 dummies with and without the new anchoring structures in the design.

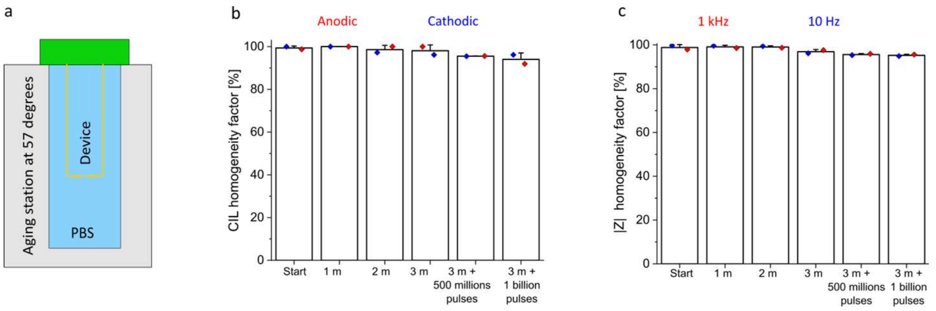


**Figure S6 | Homogeneity factor during stress test (accelerated ageing and long-term stimulation)**. **a,** Schematic of the ageing set-up. A functional device is immersed in a beaker with 10 mM PBS in a controlled temperature station at 57 °C. **b,** Homogeneity factor, for the cathodic and anodic charge injection limits, after every month of soaking in PBS at 57 °C (n=8), and after every 500 million pulses (n=4). **c,** Homogeneity factor for the impedance magnitude at 1 kHz and 10 Hz, for the conditions of subpanel b.

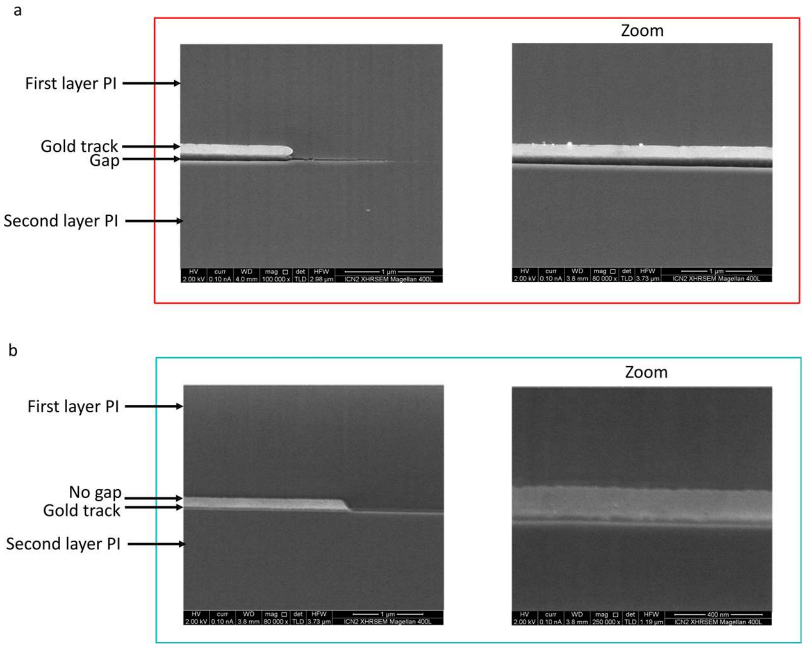


**Figure S7 | Cross-section SEM to assess interlayer adhesion. a**, SEM picture after FIB, to evaluate the cross section of a device without alumina, after 1 week soaking in PBS at 57° C. **b**, the cross section of a device with the hybrid encapsulation with alumina, after 3 months soaking.

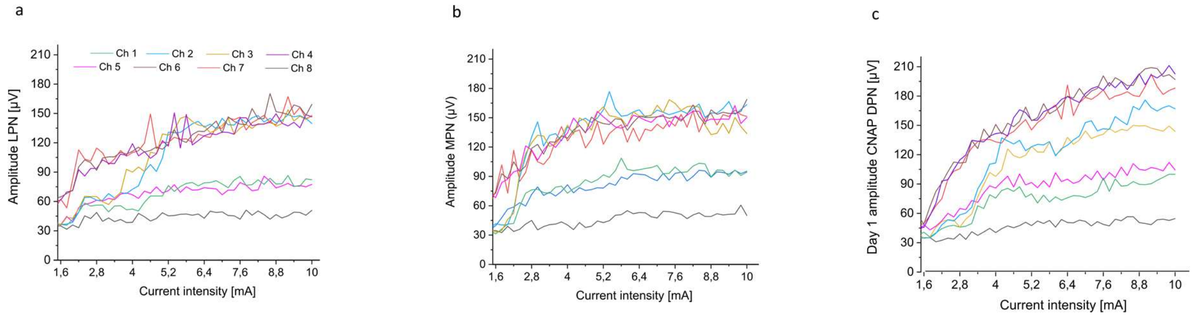


**Figure S8| Recruitment curves of the CNAPs a-c**, Recruitment curves of the peak-to-peak amplitude of the CNAPs recorded with all the electrodes of one array (n=8), elicited by stimulating with needles the LPN, the MPN, and the DPN, with biphasic current pulses (100 µs per phase, 3 Hz) at increasing current intensity.

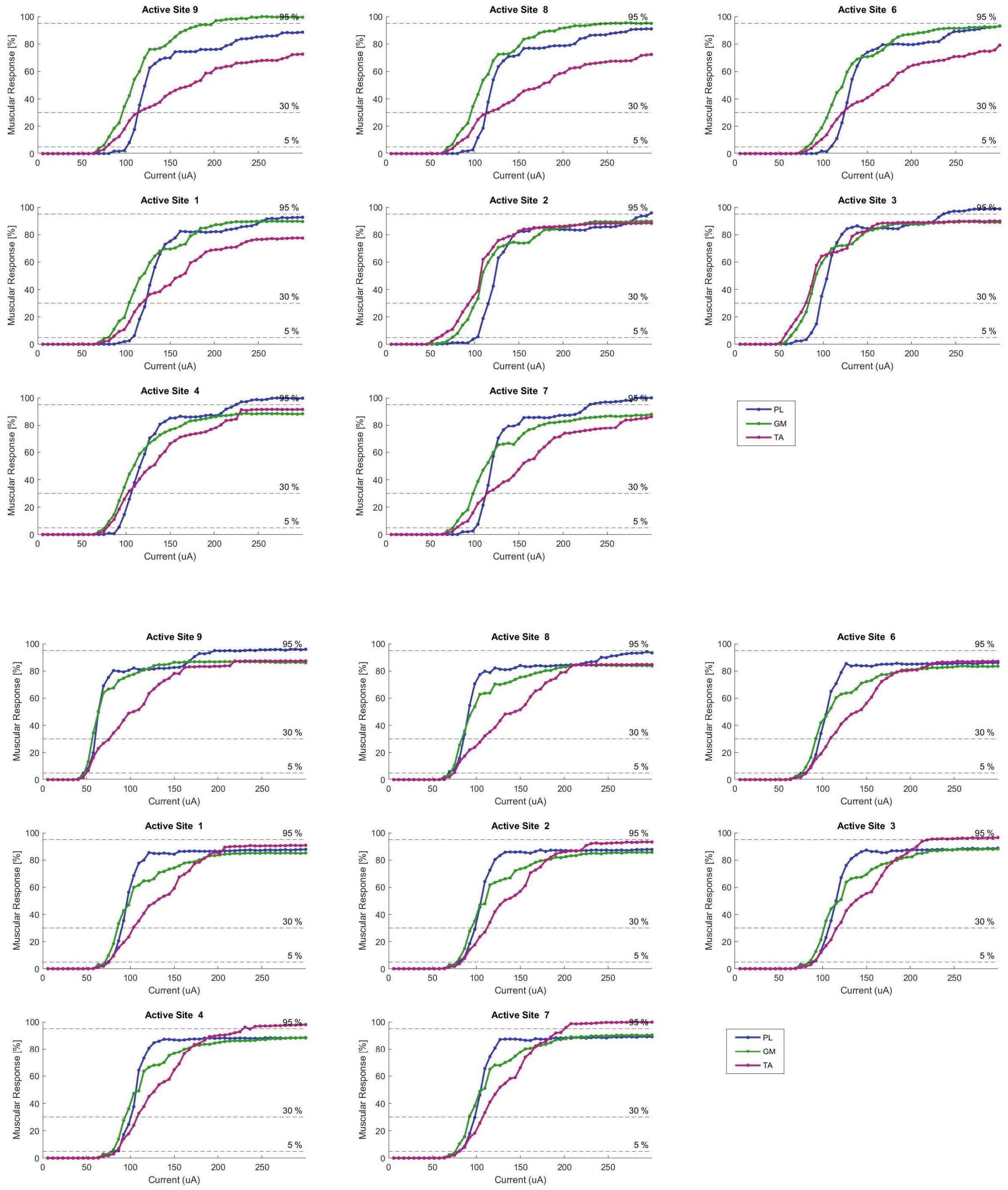


**Figure S9 | Recruitment curves after 120 days of implantation. a,** Percentage of plantar (blue), gastrocnemius (green), and tibialis anterior (red) muscle activation, during stimulation with the 8 electrodes of the 2 arrays of one device (array A on top, and array B below). Biphasic current pulses (50 µs per phase) at increasing current amplitude were injected after 120 days of implantation. The dashed lines indicate the 5, 30, and 95% of the muscle activation.

.

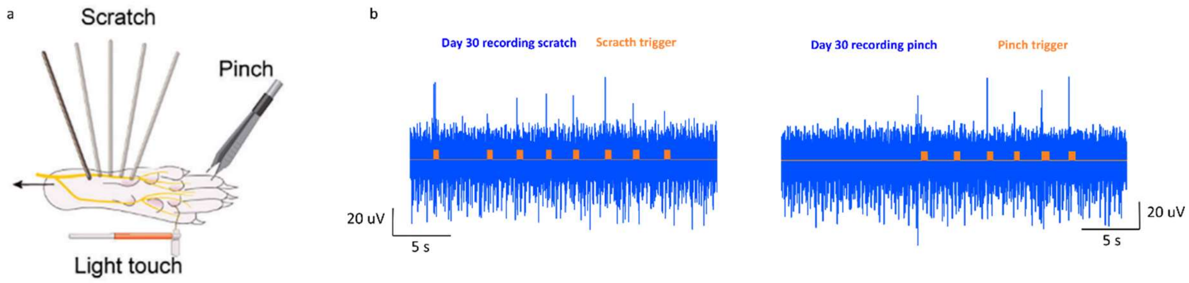


**Figure S10 | Neural recordings in chronic implantation. a,** Schematic of the experimental set-up to scratch, pinch and light touch the paw of a rat. **b**, Representative recording (light blue) in the sciatic nerve when the paw is scratched and pinched after 30 days of implantation.  The trigger (orange) indicates the time when the stimulus is performed.

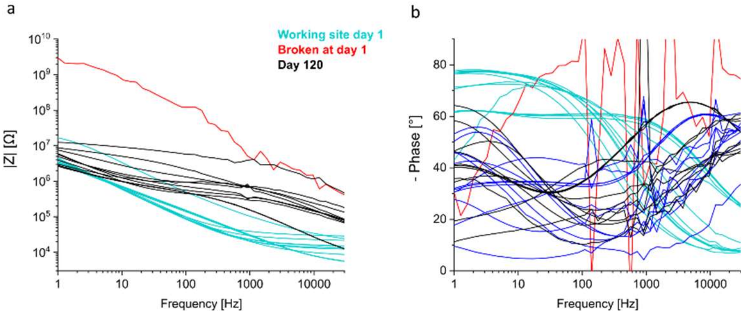


**Figure S11 | In vivo impedance over time. a,** In vivo impedance magnitude measured with one array of electrodes at day 0 (light blue), and after 120 days (black). The red trace indicates the impedance of a broken electrode at day 0 as a reference. **b,** Phase for the plots of subpanel a

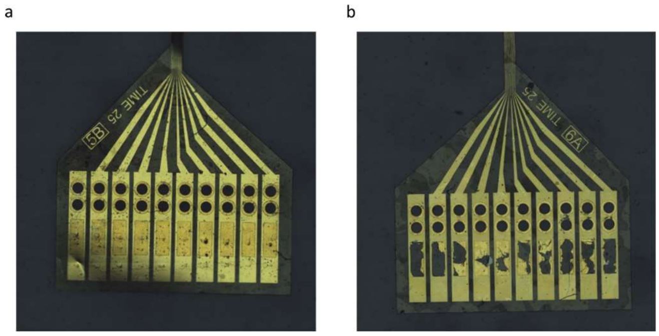


**Figure S12 | Pad area after explantation**.  **a**, Images of the pads area for a device with alumina after 1 month of implantation. **b**, Image of the pads area for a device without alumina after 1 month of implantation.

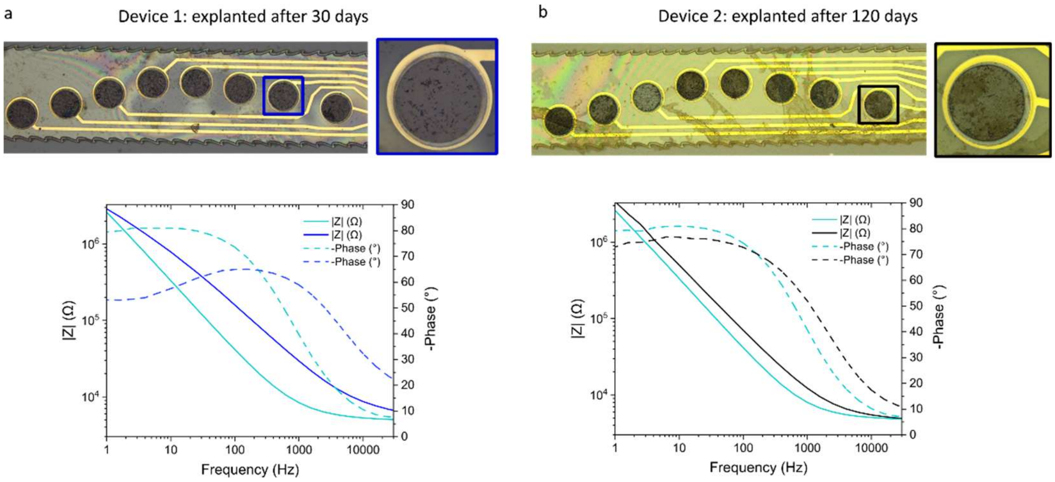


**Figure S13 | Functionality after explantation. a,** (Top) Optical image of an array explanted after 30 days, with a magnified view of one electrode. (Below) Average impedance magnitude (continuous line) and phase (dashed line) of the electrode before (light blue) implantation and after explantation (blue). **b**,( Top) Optical image of an array explanted after 120 days, with a magnified view of one electrode. (Below) Average impedance magnitude (continuous line) and phase (dashed line) of the electrode before (light blue) implantation and after explantation (black).

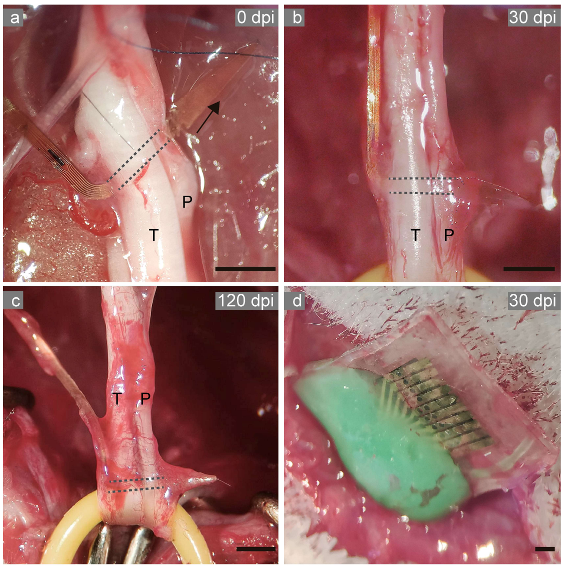


**Figure S14 Implantation procedure | a–c,** Representative images of the TIME device implanted transversally into the tibial (T) and peroneal (P) branches of the sciatic nerve. The tentative path of the TIME within the nerve is indicated by dashed lines. After implantation, fibrotic tissue covers the device, as observed in panels b and c. **d,** A plastic envelope is used to protect the pad area of the TIME device, placed subcutaneously. Scale bar: 1 mm.
